# Rearing *Istocheta aldrichi* (Diptera: Tachinidae) from field-collected Japanese beetles (*Popillia japonica*): 1. Methods to improve insect collection and parasitoid pupariation

**DOI:** 10.64898/2026.02.18.706618

**Authors:** Simon Legault, Josée Doyon, Paul K. Abram, Jacques Brodeur

**Affiliations:** Université de Montréal, Institut de recherche en biologie végétale, Département de sciences biologiques, Montréal, QC, Canada; Agriculture and Agri-Food Canada, Agassiz Research and Development Centre, Agassiz, BC, Canada

**Keywords:** Biological control, parasitoid, modified traps, insect pest, life history, Scarabaeidae, Tachinidae

## Abstract

*Istocheta aldrichi* (Diptera: Tachinidae), a specialist parasitoid of the invasive Japanese beetle, *Popillia japonica* (Coleoptera: Scarabaeidae), was released to eastern North America in the 1920s as part of a classical biological control program. Further releases are being considered in different regions of North America and Europe where *P. japonica* is establishing. Successful releases of the biocontrol agent depend on identifying efficient techniques for collecting parasitized hosts from the field and rearing the parasitoid through diapause to obtain *I. aldrichi* adults. In this study, we evaluated how the collection date, the collection method (hand-picking *vs.* regular traps vs. modified traps) and rearing conditions (food provision and substrate type) of parasitized hosts influence *I. aldrichi* pupariation and emergence. The proportion of parasitized beetles yielding *I. aldrichi* puparia decreased considerably as the season progressed. Rearing conditions immediately after collection influenced both puparium yield and quality: withholding food from parasitized *P. japonica* slightly increased puparium yield but reduced puparium weight, while the effect of food provision on subsequent overwintering survival depended on rearing substrate. Finally, simple modifications to commercial traps (larger, ventilated, containers with added food source and substrate) collected more beetles than regular traps and promoted successful development of the parasitoid to the puparium stage. Our results are used to suggest basic guidelines for collecting and rearing *I. aldrichi* in experimental research and applied biological control of *P. japonica*.

## Introduction

An important step toward establishing successful classical (importation) biological control programs is to identify efficient collection and rearing techniques once a natural enemy has been selected for release (Van Driesche & Bellows, 1993; Sheppard et al., 2019). Successful establishment of a candidate biological control agent in a new location requires the capacity to produce large numbers of high-quality individuals for release.

Methods are usually developed to mass rear agents in quarantine or laboratory facilities in such a way that they can be stored in a specific life stage and accumulated for subsequent releases. However, in cases when mass rearing a potential parasitoid agent and/or its host, is labor intensive and too costly, parasitized hosts can be collected directly from wild populations. This approach has the advantage of preventing the decrease of the intrinsic genetic diversity and fitness-related life-history traits often associated with continuous rearing for extended periods (Hopper et al., 1993; Leung et al., 2020).

The Japanese beetle, *Popillia japonica* Newman (Coleoptera: Scarabaeidae), first invaded eastern North America in the early 1900s where it spread rapidly and became a serious pest of a wide range of ornamentals and field crops (Althoff & Rice, 2022). More recently, new populations have been detected in several parts of the world, including Western North America and Europe (Strangi et al. 2025; Brodeur et al. 2024), causing a renewed interest for classical biological control in these regions (CABI, 2021; Abram et al., 2022; Makovetski & Abram, 2024).

More than a century ago, *Istocheta aldrichi* (Mesnil) (Diptera: Tachinidae) has been identified as the most promising classical biological control agents for *P. japonica* (Clausen et al., 1927), its only known host based on existing documented Tachinidae-host associations (Makovetski et al., 2025). Parasitoids were hand-collected from parasitized *P. japonica* in Japan, brought to a United States Department of Agriculture laboratory, and adults were first released in 1922 near Moorestown, NJ, USA. From 1923 to 1950, the parasitoid was redistributed in 12 other northeastern US states (Fleming, 1968). More recently, additional redistributions have been conducted in Minnesota, Colorado and North Carolina (McDonald & Klein, 2023). Meanwhile, the parasitoid likely spread northwards naturally, and became established in the Canadian provinces of Ontario and Québec (O’Hara, 2014; Gagnon et al., 2023). In 2023-2025, *I. aldrichi* was successfully redistributed from Québec and Ontario to British Columbia (Makovetski & Abram, 2024) in order to supplement other tactics (mostly insecticide applications) being used to eradicate *P. japonica* populations. Redistributions are currently planned in other Canadian provinces and introductions are being considered in Europe (CABI, 2021).

In northeastern North America, *I. aldrichi* adults are mostly active at the beginning of the summer, before the peak emergence of *P. japonica* adults (King, 1931; Gagnon et al., 2023). Female *I. aldrichi* typically lay one large, conspicuous, white egg on the pronotum of *P. japonica* adults (Clausen et al., 1927; Pelletier et al., 2023). Upon eclosion, *I. aldrichi* larva drills downwards into the beetle and feeds on host tissues. Parasitized *P. japonica* next burrow into the soil and die. The *I. aldrichi* larvae pupariate within the host cadaver, overwinter, and the adults emerge in early summer of the following year. Parasitism levels of *P. japonica* then decrease rapidly from late July to early September, while *P. japonica* remain abundant (Gagnon et al., 2023). There is only one generation per year (Clausen et al., 1927).

Despite the long history of biological control programs using *I. aldrichi*, methods to collect and rear this biological control agent need to be optimized. Previous introduction and redistribution programs have relied on collecting parasitized *P. japonica* from natural populations in Japan or established populations in new regions (Clausen et al., 1927; Fleming, 1968; McDonald & Klein, 2023; Makovetski & Abram, 2024), using either hand collections or Japanese beetle traps. Hand collection of parasitized *P. japonica* can be efficient when parasitism levels are high (over 15–20%) but are otherwise time-consuming. Furthermore, *P. japonica* adults easily drop to the ground or fly away when startled (Kreuger & Potter, 2001), making it challenging for inexperienced workers to collect large numbers of parasitized hosts. On the other hand, highly efficient traps baited with floral compounds and a sex pheromone (Tumlinson et al., 1977; Ladd & Klein, 1986) have long been commercialized. These traps equally attract parasitized and unparasitized adult beetles of both sexes, with no particular sampling bias regarding the body size of trapped individuals (Legault et al., 2024). As a result, these traps are suitable for mass collections of *I. aldrichi* individuals that are representative of a natural population. Unfortunately, our preliminary observations indicate that captured individuals generally die rapidly of heat and/or drowning in the trap recipient, resulting in very low pupariation rate of *I. aldrichi*.

Following field collection of parasitized beetles, developing parasitoids must be maintained in suitable rearing conditions to reach the puparium stage, overwinter, and emerge as adults the following year. In previous attempts to rear parasitized beetles, plant material was supplied on top of a substrate (e.g., soil, vermiculite, sphagnum moss) to both feed the beetles and provide them with a substrate in which they can bury themselves before they die and the parasitoid larva forms a puparium (Clausen et al., 1927; Fleming, 1968; Kidd & McDonald, 1992; McDonald & Klein, 2023). Variations on these methods (e.g. using different substrates) have not been formally trialed in controlled experiments, however.

To produce *I. aldrichi* for redistribution, more knowledge is needed to best collect parasitized *P. japonica*, rear parasitoids through critical events in their seasonal life cycle (e.g., larval and pupal development, diapause), schedule their emergence for biological control releases, and maximize adult survival and early egg production after emergence and before release. The present study examines aspects related to field collection of parasitized beetles and further rearing the parasitoid to the puparium stage. The effect of various factors on the downstream stages of *I. aldrichi* rearing – i.e., how parasitoid puparia are stored and overwintered, and how emergence of adults can be timed with host availability the next season – were tested in a companion study (Abram et al., 2026).

In this study, we asked if simple modifications to the recipient of commercialized Japanese beetle traps, mainly increased ventilation and rain draining, would maintain parasitized *P. japonica* alive long enough for the successful development of *I. aldrichi* into puparia within traps. Because *I. aldrichi* kills its host rapidly (Clausen et al., 1927; Clausen, 1956), we further tested if providing food to trapped parasitized beetles would influence *I. aldrichi* pupariation rate, puparium weight, and overwintering survival. We also experimentally tested the importance of capturing parasitized beetles early during the activity period of *I. aldrichi* to increase parasitoid pupariation rate.

## Methods

### Experiment #1: Effect of collection date on I. aldrichi pupariation

To assess the effect of collection date on *I. aldrichi* pupariation success, parasitized *P. japonica* adults were collected in 2024 in a commercial vineyard (Coteau St-Paul) located in the municipality of Saint-Paul-Abbotsford, Québec, Canada (Site 1). *Popillia japonica* adults bearing one or multiple *I. aldrichi* egg(s) on their pronotum were haphazardly hand-picked on common grape vine (*Vitis vinifera*) about 3-5 times per week during the period for which they were present on grape vines (i.e., from July 3 to August 15). Beetles were collected on sunny days, between 9:00 and 14:00, which is when adults are the most active on the top of host plants (Vittum, 1986; Legault et al., 2024). Parasitized beetles were approached slowly to avoid startling them and gently caught between the thumb and forefinger of an experienced worker. Beetles were placed in a container with fresh grape vine leaves and brought back to the laboratory. They were then placed in cohorts of up to 230 individuals (1-3 cohorts per collection date) in 1L ventilated plastic containers with moistened vermiculite for substrate (Figure 1A). Containers were stored in a controlled environment (21±1°C, 45±5% RH, 14L:10D), and parasitized beetles were fed *ad libitum* with fresh grape vine leaves for two weeks. After that period, the content of each container (cohort) was spread on a sorting tray and carefully examined to retrieve parasitized *P. japonica* and loose *I. aldrichi* puparia. Beetles were dissected under a stereomicroscope to extract all remaining *I. aldrichi* puparia. For each cohort, *I. aldrichi* pupariation rates were determined by dividing the number of *I. aldrichi* puparia by the number of parasitized beetles collected and reared (N= 8–230).

**Figure 1.**
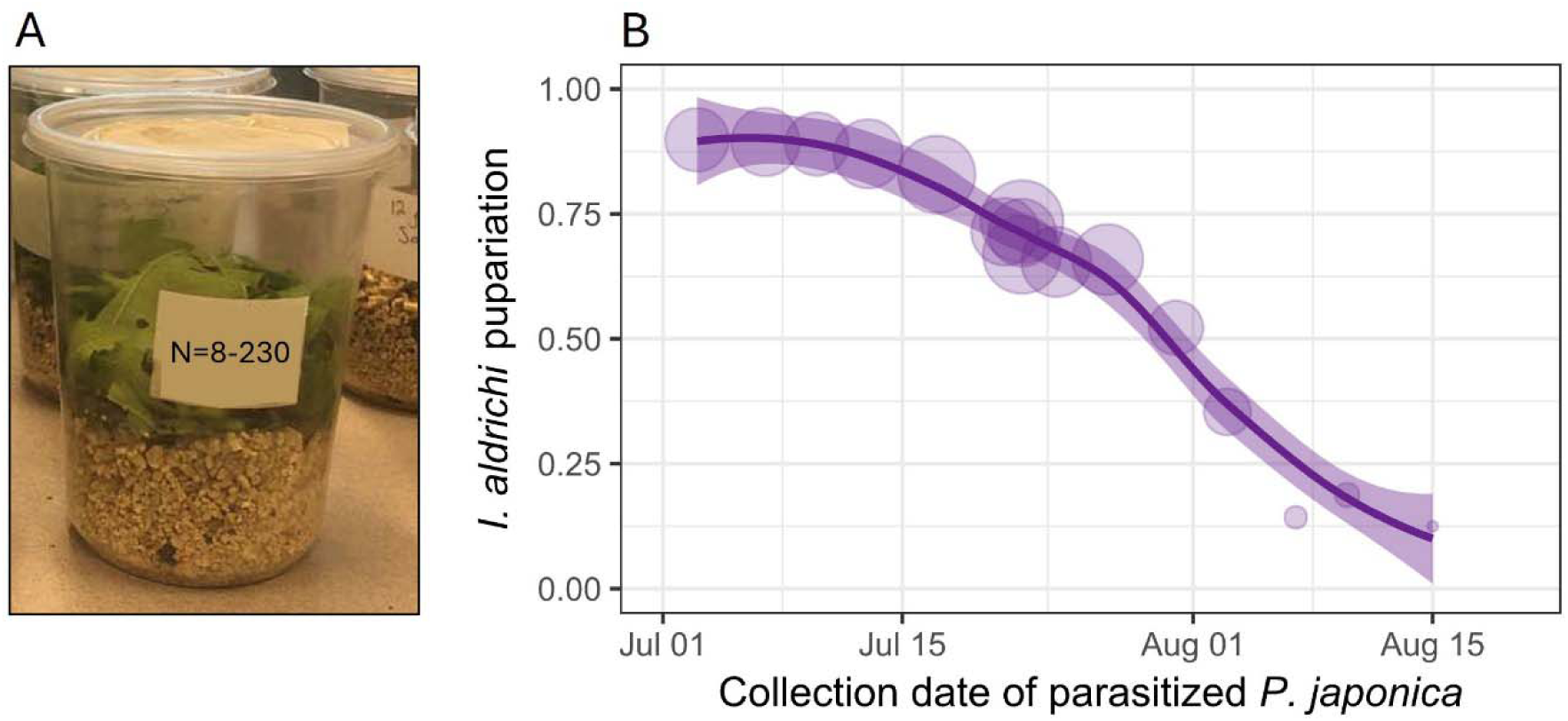
**A)** Ventilated plastic containers with moistened vermiculite for substrate and fresh grape vine leaves to feed *P. japonica* parasitized by *I. aldrichi*. Photo by S. Legault. **B)** *Istocheta aldrichi* proportion pupariation as a function of collection date of parasitized *P. japonica*. The size of dots is proportional to the number of reared beetles (N=8–230).

### Experiment #2: Effects of P. japonica feeding and substrate on I. aldrichi pupariation and adult emergence

The goal of this factorial experiment was to determine whether providing field-collected parasitized *P. japonica* with food and varying their rearing substrate affects the yield and size of *I. aldrichi* puparia, and the parasitoid overwintering survival. *Popillia japonica* parasitized by *I. aldrichi* were haphazardly hand-picked on grape vines on July 11 and July 18, 2023, between 9 and 11 AM in Site 1. Parasitized hosts were placed in a container with fresh grape vine leaves and brought back to the laboratory. They were then placed in cohorts of 100 individuals in 1 L, ventilated, plastic containers with either moistened vermiculite or all-purpose garden soil (VALU+, RONA) for substrate (Figure 2A). Beetles were either fed *ad libitum* with fresh grape vine leaves or deprived of food (N=6 cohorts/repetitions per combination of factors). Rearing containers were placed in a controlled environment (21±1°C, 45±5% RH, 14L:10D) for 20 days. After that period, *I. aldrichi* puparia were retrieved as in Experiment 1. For each cohort, the proportion of parasitized *P. japonica* yielding *I. aldrichi* puparia was determined by dividing the number of parasitoid puparia by 100. Average parasitoid puparium weights per treatment were determined by weighing together all puparia of each cohort and dividing the values by the number of puparia. Puparia obtained from each cohort were individually stored in 60 ml plastic cups filled with 0.5 cm of moist vermiculite for substrate in a controlled environment (21±1°C, 45±5% RH, 14L:10D) until October 1, 2023. Puparia were then shipped to the Agassiz, Research & Development Centre, of Agriculture and Agri-Food Canada in British Columbia on November 14, 2023, and each sample was transferred to plastic ventilated containers (9.5 cm × 6 cm × 6 cm), and overwintered in a walk-in growth chamber set to 5°C. Temperature settings were monitored with a temperature logger taking readings every 30 min (actual average: 5.43°C). On April 21, 2024, containers with puparia were transferred to a growth chamber set at 1.5°C lower than the previous week’s average soil temperature in Agassiz, BC (Table S1), and the substrates were moistened with water as needed. Adult parasitoids from each container were counted as they emerged, and proportion emergence was calculated as the number of emerging adult flies divided by the total number of puparia in each sample.

**Figure 2.**
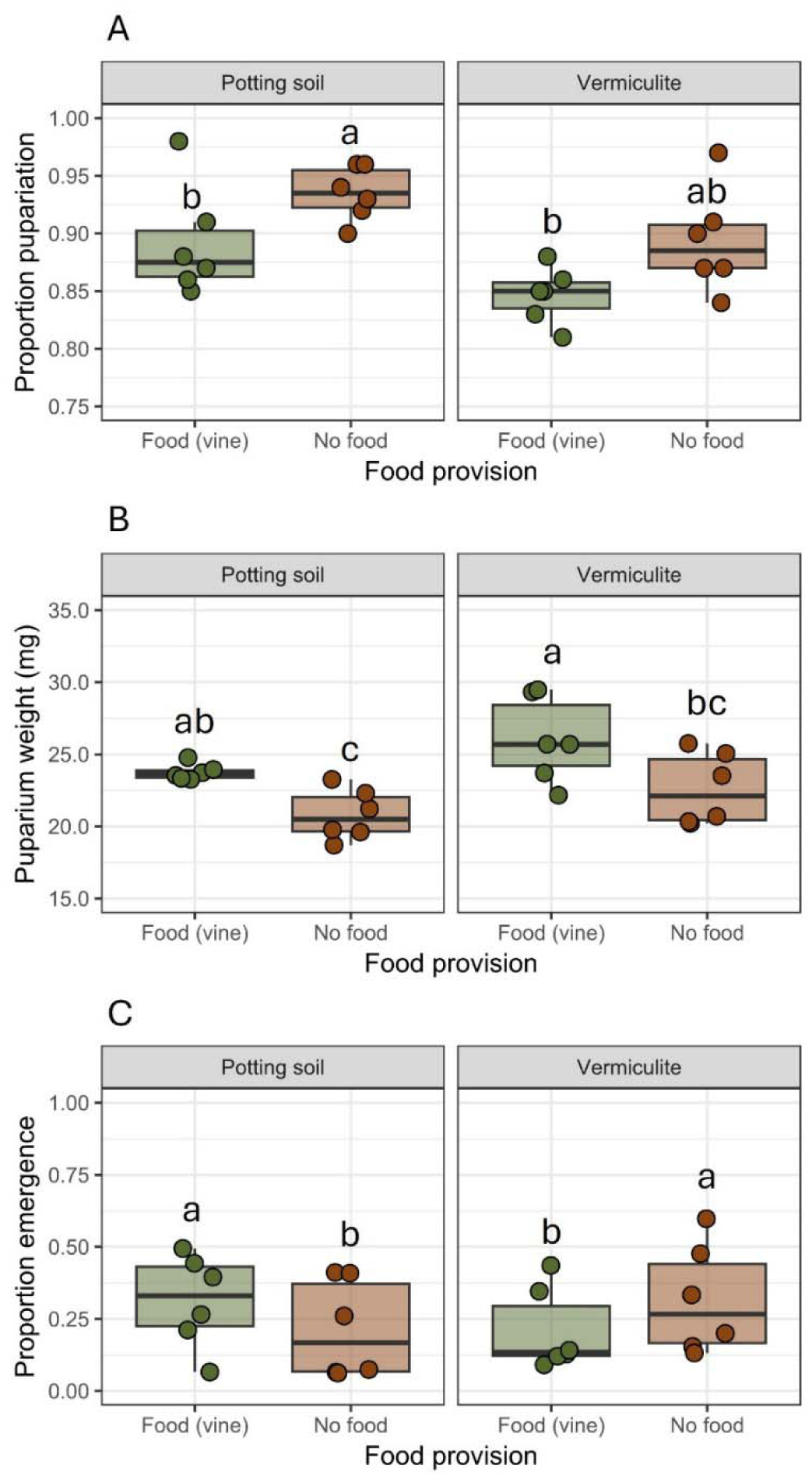
**A)** *Istocheta aldrichi* proportion pupariation, **B)** mean puparium weight, and **C)** proportion adult emergence after overwintering. Each point represents a cohort of 100 *P. japonica* adults parasitized by *I. aldrichi* reared in containers with a layer of potting soil or vermiculite as substrate and fed *ad libitum* with fresh grape vine leaves or deprived of food. Different letters above boxes indicate significant differences between treatments.

### Experiment #3: Effects of trap modifications on P. japonica abundance and I. aldrichi pupariation and adult emergence

To maximize trap capture capacity and pupariation rate of *I. aldrichi*, the customary 1.5L container of Bioprotec© traps (Figure 3A) was replaced by a 19 L white plastic bucket, pierced on the sides with four ventilation holes (5cm diameter, at mid height of the buckets) covered with muslin (Figure 3B). About 30 small 1.5 mm holes were also drilled in the bottom of the container to drain excess water. (Figure 3B).

**Figure 3.**
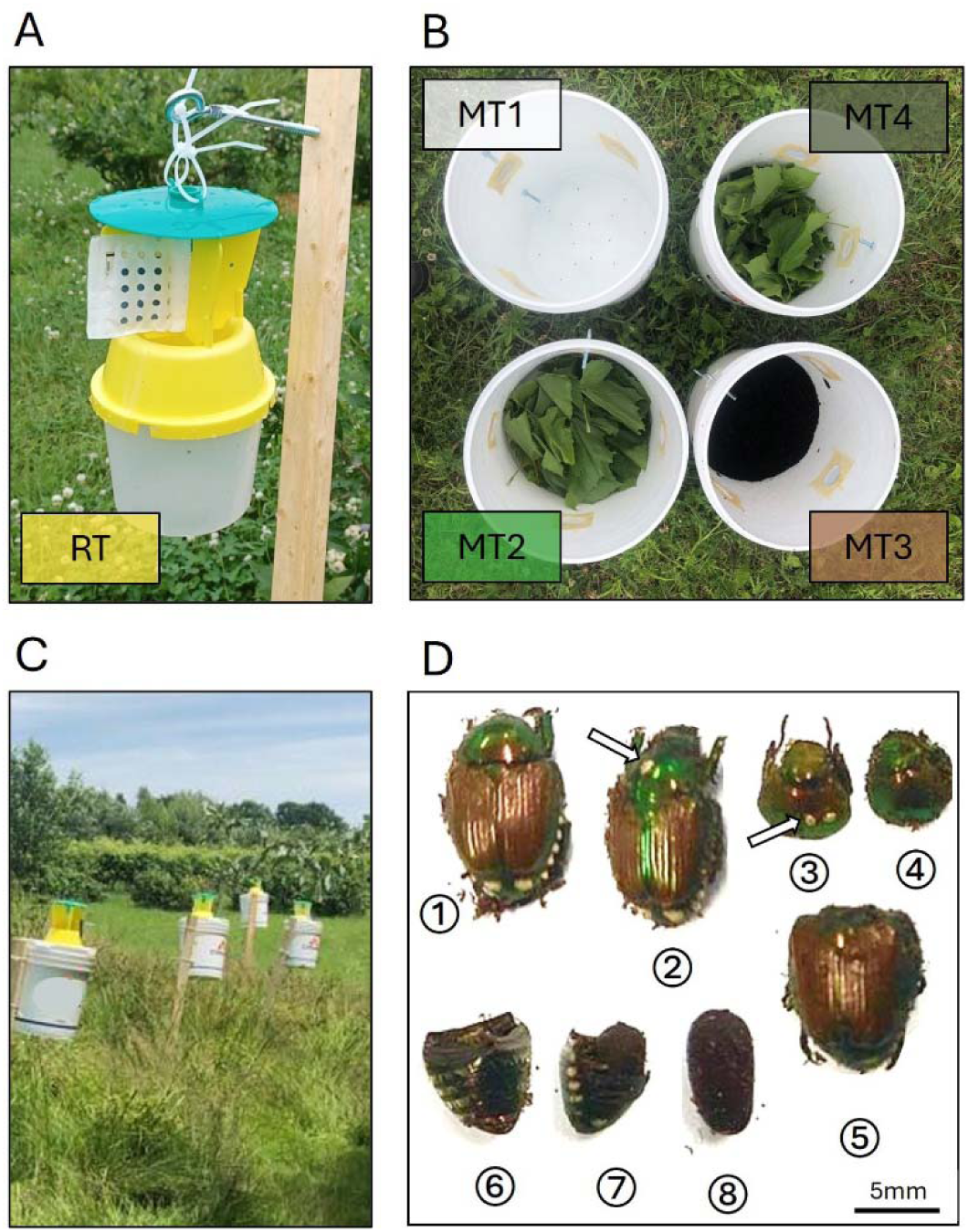
**A)** A regular Bioprotec© trap (RT) installed at the top of a 1m wooden stake. Photo by C. Lacroix. **B)** Content of the modified traps: empty (MT1), grape vine leaves (MT2), black earth (MT3), grape vine leaves and black earth (MT4). Photo by S. Legault. **C)** Traps placed along a linear transect (replicated block). Photo by S. Legault. **D)** Insects and insect parts found in Japanese beetle traps: ⍰ complete, unparasitized *P. japonica*, ⍰ complete, parasitized *P. japonica*, ⍰ loose head and thorax of a parasitized beetle ⍰ loose head and thorax of an unparasitized beetle, ⍰ loose *P. japonica* abdomen, ⍰ loose empty *P. japonica* abdomen, ⍰ loose *P. japonica* abdomen containing an *I. aldrichi* puparium, ⍰ loose *I. aldrichi* puparium. *Istocheta aldrichi* eggs are indicated with arrows. Photo from S. Legault.

Field trials were conducted in mid-July 2024, when parasitized *P. japonica* are the most abundant in southern Québec, Canada (Gagnon et al., 2023). Traps were installed at Site 1 and at a second vineyard (Vignoble du PicBois) with an history of *P. japonica* infestation located in the municipality of Brigham, Québec, Canada (Site 2). Spatially replicated blocks were deployed on July 10 (Site 1) and July 11 (Site 2) and consisted of five different trap treatments: a regular trap (RT) as a control, and modified traps described above (MT1-MT4). To assess the impact of substrate and food availability on *I. aldrichi* pupariation rate, we added to modified trap containers either: nothing (MT1), fresh, pesticide-free, grape vine leaves (MT2), black soil for outdoor gardening (VALU+, RONA; MT3) and grape vine leaves and black soil (MT4) (Figures 3A and B). All traps were installed at the top of a 1 m wooden stake along a linear transect of approximately 25 m (5 m between traps; Figure 3C). The relative position of each trap treatment was determined randomly for each block. We installed four replicated blocks per site, each separated by at least 100 m. Trapping was run for seven days to avoid trap saturation, for two consecutive sampling periods (2 sites × 4 replicated blocs × 2 trapping periods = 16 trap samples per treatment). At the beginning of each trapping period, about 2L of commercial black soil were added to the containers of MT3 and MT4, together with four handfuls of fresh grape vine leaves to MT2 and MT4. Because food was quickly consumed by trapped beetles, it was refreshened with the same quantity of grape vine leaves at day 4. After each trapping period, traps contents, including substrate and leaves for MT2-MT4, were transferred into a 3.5 L plastic container with ventilated lids. Containers were brough back to the laboratory and stored in darkness (21±1°C, 45±5% RH, 0L: 24D) for two weeks to allow *I. aldrichi* to pupariate.

The content of each trap was next meticulously examined under a stereomicroscope to count the following insects and insect parts: unparasitized and parasitized *P. japonica* (see ⍰ and ⍰, respectively; Figure **3**D), loose unparasitized (⍰) and parasitized heads and thoraxes (⍰), loose abdomens (⍰, ⍰ and ⍰), and loose *I. aldrichi* puparia (⍰) (Figure **3**D). *Popillia japonica* bearing *I. aldrichi* eggs as well as loose beetle abdomens were dissected to retrieve all *I. aldrichi* puparia. For each trap sample, we determined the: i) total number of trapped *P. japonica* by counting the total number of abdomens (⍰+⍰+⍰+⍰+⍰), ii) proportion parasitism by dividing the number of thoraxes bearing at least one parasitoid egg and the total number of thoraxes ([⍰+⍰]/[⍰+⍰+⍰+⍰]), and iii) *I. aldrichi* pupariation rate, by dividing the number of *I. aldrichi* puparia by the number of parasitized beetles.

After sorting, *I. aldrichi* puparia obtained from each trap sample were individually stored in 60 ml plastic cups filled with 0.5 cm of moist vermiculite until September 24, 2024 (i.e., about 65 days post-collection). Puparia were then transferred to small muslin bags (3 × 4 cm) and buried outdoors in a lawn plot (20 cm deep) at the Montréal Botanical Garden, Canada. Muslin bags were covered with chicken wire (gauge galvanized steel) for protection against predation by small animals. Overwintered puparia were recovered on June 2, 2025 (i.e., after ∼8.5 months). Up to 50 puparia from each trap sample were used to measure adult parasitoid emergence rates (remaining puparia were used for other experiments not described in this study). Puparia were transferred to ventilated insect rearing boxes (ProLab Supply, W7 × D7 × H10 cm) without the lids and stored in cool and dark conditions (12 ± 1 °C, 50 ± 5% RH, 0L: 24D) for 23 days to delay fly development in prevision of field releases for another study. This procedure does not reduce emergence success (Abram et al., 2026). Boxes were next transferred to BUGDORM® insect rearing cages (W33 × D33 × H77 cm) and placed in warmer conditions (22 ± 1 °C, 80 ± 5% RH, 16L: 8D) to promote parasitoid development. Cages were monitored daily for parasitoid emergence and adults were captured and sexed based on the presence (females) or absence (males) of two proclinate outer orbital setae (J. O’Hara, personal communication). After the fly emergence period, puparia from each trap sample/rearing box were examined individually under a stereomicroscope to measure fly emergence rates by dividing the number of puparia opened by emerging flies (i.e., with a characteristic circular opening), by the total number of examined puparia (i.e., up to 50 puparia from each trap sample).

Voucher specimens of *I. aldrichi* adults from this study have been incorporated into the Canadian National Collection of Arthropods and Nematodes (Ottawa, Ontario).

## Statistical analyses

For Experiment 1, a generalized linear model (GLM) with a binomial error distribution was used to assess the effect of collection date on *I. aldrichi* proportion pupariation. Sample sizes for each *P. japonica* cohort were used as prior weights in the fitting process. The statistical significance of collection date was assessed with Wald z-statistic.

For Experiment 2, we used generalized linear mixed models (GLMM) with a binomial error distribution to assess the effects of food provision, substrate type, and their interaction, on proportion pupariation and overwintering survival of *I. aldrichi*. We used a linear mixed model (LMM) to assess the effects of these factors on puparium weight. For each model, sampling dates were included as random variables. The statistical significance of each model term was assessed with type II Wald chi-squared (GLMM) or F-tests (LMM).

For Experiment 3, we used generalized linear mixed models (GLMM) with binomial error structures to analyse the effects of trap types on i) *P. japonica* relative abundance (number of *P. japonica* captured in a trap / total number of *P. japonica* captured in the five trap treatments of a replicated block), ii) proportion parasitism, iii) *I. aldrichi* proportion pupariation, and iv) *I. aldrichi* emergence. Sample sizes were used as prior weights in the fitting process. The statistical significance of trap treatments was assessed with type II Wald chi-squared tests. Site, replicated block, and sampling period were included as nested random variables in each model.

For Experiments 2 and 3, the significance of pairwise differences between groups were assessed with the R package *emmeans* (Lenth, 2021) using the Tukey *P* value adjustment method.

## Results

### Experiment #1: Effect of collection date on I. aldrichi pupariation

A total of 4,295 parasitized *P. japonica* was hand-collected and reared in the laboratory. The collection date strongly influenced *I. aldrichi* successful development (GLM: *z*=-12.88, *P*<0.001; Figure 1B). Pupariation rates averaged 86.4% for parasitized beetles collected between July 3 and July 17, and next gradually decreased to reach 15.2% for those collected between August 7 and August 15. No parasitized beetles were found at Site 1 after August 15.

### Experiment #2: Effects of P. japonica feeding and substrate on I. aldrichi pupariation and adult emergence

Host feeding and rearing substrates significantly influenced the proportion of parasitized *P. japonica* yielding *I. aldrichi* puparia (Table 1; Figure 2A). Parasitoid pupariation was slightly (∼4%) higher when parasitized beetles were deprived of food (estimated marginal means of 89 and 93% for vermiculite and potting soil, respectively), than when they were provided with foliage (85 and 89%). Pupariation was also slightly (∼4%) higher when beetles were reared on soil rather than vermiculite (Table 1; Figure 2A). The weight of *I. aldrichi* puparia also varied significantly with both host feeding and rearing substrate (Table 1, Figure 2B). Puparia from fed beetles were 14–15% heavier than those from unfed beetles, and puparia from beetles reared on vermiculite were 8–9% heavier than those reared on potting soil. Following overwintering, adult parasitoid emergence depended on an interaction between host feeding and rearing substrate, with the effect of food provision differing between substrates (Table 1, Figure 2C). Parasitoids reared from fed beetles were slightly more likely to emerge as adults than those from starved beetles in potting soil, while the opposite was true in vermiculite.

**Table 1.**
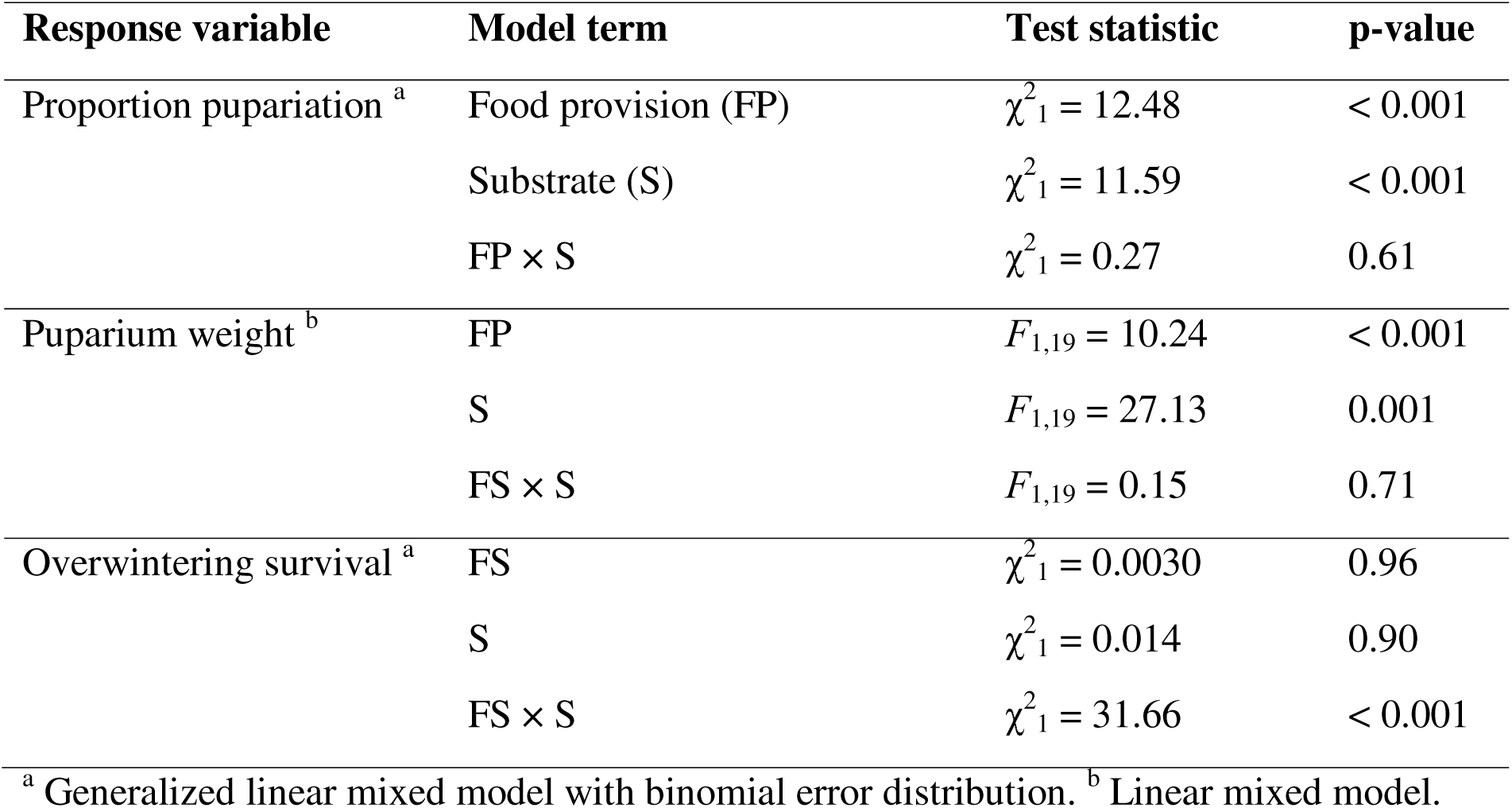
Statistical information for Experiment 2, which tested the effects of food provision (FP) and substrate (S) on proportion pupariation, puparium weight, and proportion emergence of *Istocheta aldrichi*. Information for random effect term (collection date; see Methods) is not shown. See Figure 2 for visual representations of the data and the significances of pairwise differences between combination of factors.

| Response variable | Model term | Test statistic | p-value |
| --- | --- | --- | --- |
| Proportion pupariation <sup>a</sup> | Food provision (FP) | $\chi^2_1 = 12.48$ | < 0.001 |
| | Substrate (S) | $\chi^2_1 = 11.59$ | < 0.001 |
| | FP × S | $\chi^2_1 = 0.27$ | 0.61 |
| Puparium weight <sup>b</sup> | FP | $F_{1,19} = 10.24$ | < 0.001 |
| | S | $F_{1,19} = 27.13$ | 0.001 |
| | FS × S | $F_{1,19} = 0.15$ | 0.71 |
| Overwintering survival <sup>a</sup> | FS | $\chi^2_1 = 0.0030$ | 0.96 |
| | S | $\chi^2_1 = 0.014$ | 0.90 |
| | FS × S | $\chi^2_1 = 31.66$ | < 0.001 |

<sup>a</sup> Generalized linear mixed model with binomial error distribution. <sup>b</sup> Linear mixed model.

### Experiment #3: Effects of trap modifications on P. japonica abundance and I. aldrichi pupariation and adult emergence

A total of 36,729 Japanese beetles were trapped during the experiment. All modified traps captured more beetles than regular traps, with modified traps containing grape vine foliage (MT2 and MT4) being more efficient than traps without food (MT1 and MT3) (Table 2; Figure **4A**). Modified traps with black soil only (MT3) did not capture significantly more beetles than empty modified traps (MT2). **Overall, parasitism was 24.2 ± 6.9% (mean ± SD).** We found a small, but significant effect of trap types on proportion parasitism by *I. aldrichi*, with parasitism significantly lower in MT4 compared to RT, MT1 and MT2 (Table 2; Figure **4B**). *Istocheta aldrichi* pupariation rates were strongly influenced by trap types (Table 2; Figure **4C**). Mean pupariation rates were maximum in MT4 (84.0%), followed by MT2 (67.7%) and MT3 (60.9%), and very low in MT1 (5.0%) and RT (3.8%). In three MT4 samples (out of 16), *I. aldrichi* pupariation rates exceeded 100%, suggesting an underestimation of proportion parasitism in these samples. Finally, *I. aldrichi* emergence rate was also strongly influenced by trap types (Table 2; Figure **4D**). Mean emergence rates were maximum in MT4 (20.6%), followed by MT3 (15.6%), and very low in MT2 (7.7%) RT (7.4%) and MT1 (0.0%).

**Figure 4.**
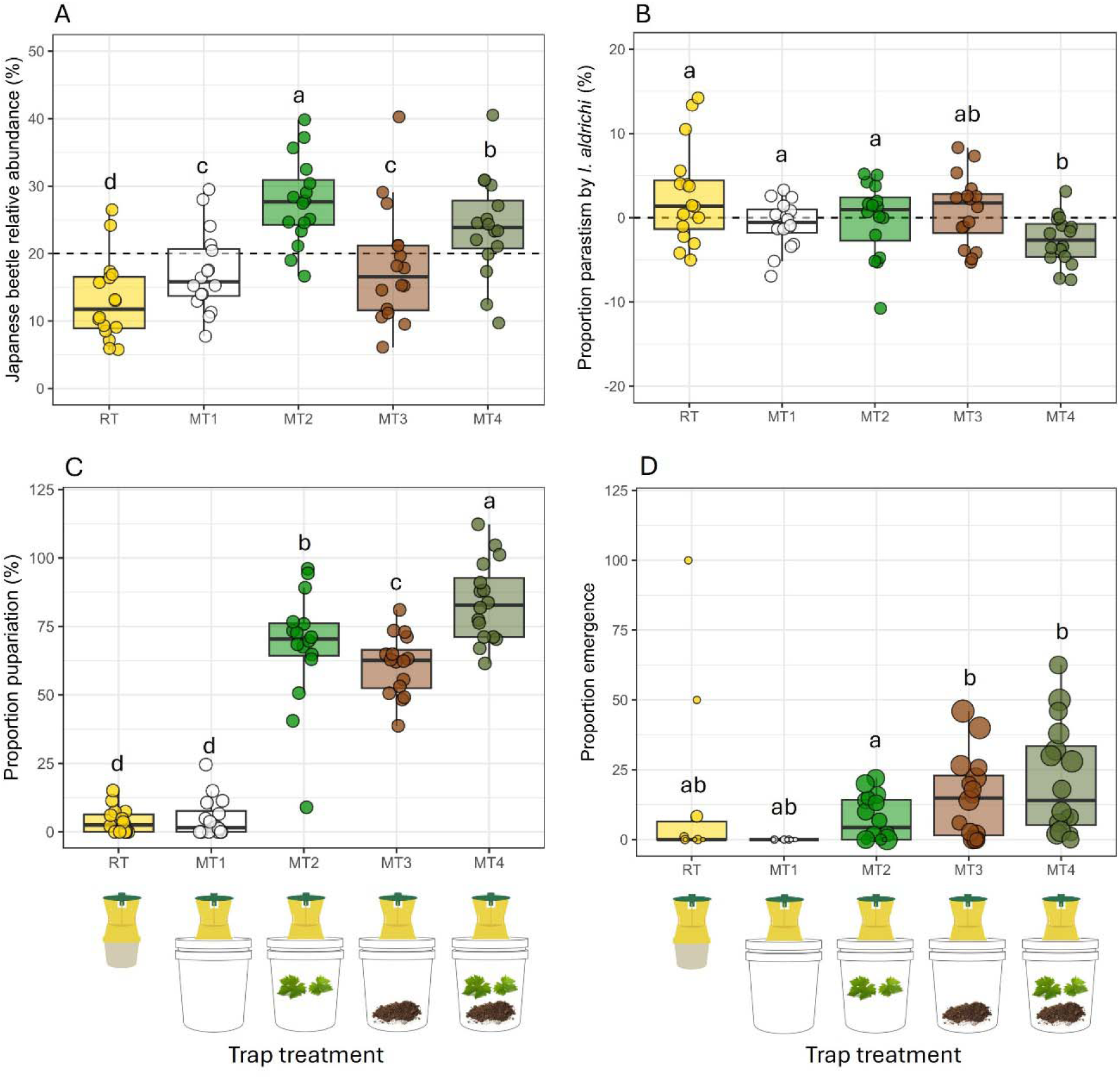
**A)** *Popillia japonica* relative abundance, **B)** proportion parasitism by *Istocheta aldrichi* (deviation from the mean within a replicate block), **C)** *I. aldrichi* proportion pupariation, and **D)** proportion emergence for the five trap treatments. Different letters above boxes indicate significant differences between trap treatments. Horizontal dashed lines indicate balanced relative abundance (20%) and equal (0% difference) proportion parasitism between trap treatments, respectively. Soil and leaves icons are from Proonty and Scisettialfio, respectively (Shutterstock.com); trap icons are from S. Legault.

**Table 2.** Statistical information for Experiment 3, which tested the effect of trap treatments (T) on *P. japonica* relative abundance, proportion parasitism by *Istocheta aldrichi*, proportion pupariation, and proportion emergence using generalized linear mixed model with binomial error distribution. Information for random effect terms (site, replicated block, and sampling period; see Methods) is not shown. See Figure 3 for visual representations of the data and the significances of pairwise differences between trap treatments.

| Response variable | Model term | Test statistic | p-value |
| --- | --- | --- | --- |
| Relative abundance | Trap treatment (T) | $\chi^2_4 = 238.52$ | < 0.001 |
| Proportion parasitism | T | $\chi^2_4 = 29.52$ | < 0.001 |
| Proportion pupariation | T | $\chi^2_4 = 1677.6$ | < 0.001 |
| Proportion emergence | T | $\chi^2_4 = 27.48$ | < 0.001 |

## Discussion

In this study, we identified several determining factors for the successful field capture of parasitized *P. japonica* and subsequent rearing of *I. aldrichi*. Because continuous rearing of *P. japonica* under laboratory conditions is both challenging (Ludwig & Fox, 1938) and costly, and because parasitized *P. japonica* can be relatively abundant in the field, the practical method to obtain *I. aldrichi* for research and biological control programs is from field collections (Clausen et al., 1927; McDonald & Klein, 2023; Makovetski & Abram, 2024). However, as we showed, successful rearing of *I. aldrichi*, from collection of parasitized *P. japonica* in the field to spring emergence of diapausing individuals, depends on a few major sources of variability. These include the collection date and method (hand-picked *vs.* regular traps *vs.* modified traps), and to a lesser degree food provisioning to parasitized beetles and the type of substrate used. We discuss these aspects below and conclude with some recommendations.

### Collection date

Parasitized *P. japonica* collected early in the season yield much more *I. aldrichi* puparia than later-collected hosts. There are at least two explanations for this observation. First, small proportions of beetles bearing a *I. aldrichi* egg survive parasitism (4.2 % of feed beetles in Experiment 2), probably because of unhatched parasitoid eggs or parasitoid larvae that died early during development (Clausen et al., 1927). Such beetles, still bearing the eggshells of *I. aldrichi*, would thus accumulate throughout the season, and can be observed as late as mid-September (Gagnon et al., 2023; Makovetski et al., 2025). As a result, the proportion of *P. japonica* bearing an *I. aldrichi* egg that produces a parasitoid puparium likely decreases when collected later in the season. Second, and to a lesser extent, later-collected hosts might also be less suitable for *I. aldrichi* development, because older hosts have been shown to be of lower quality for many parasitoid species (Vinson & Iwantsch, 1980).

### Collection method

We tested different methods to collect parasitized *P. japonica* and preserve them during parasitoid larval development. Hand-picking is a valid option but requires some experience in order to catch the startled beetles before they fly away or fall to the ground (Kreuger & Potter, 2001). Although time-consuming (about 1 hour to collect ∼200 parasitized beetles during Experiment 1), hand-picking is effective for collecting small numbers of individuals, e.g., for laboratory experiments. Alternatively, commercial traps with modified containers and added plant material and substrate are very effective at i) attracting and collecting large numbers *P. japonica* (including parasitized individuals), and ii) enhancing parasitoid pupariation rates, compared to regular traps.

Replacing the 1.5-L regular trap container with a 19-L ventilated bucket increased *P. japonica* capture rate by 57% on average. Improved ventilation rather than larger container volumes likely explains this result. Ventilation would lower temperature inside trap containers, thereby reducing decomposition rates of trapped beetles and the resulting odors that either repel *P. japonica* or mask volatile floral compounds and *P. japonica* sex pheromone of baited commercial traps (Alm et al., 1996). Piñero and Dudenhoeffer (2018) also showed that ventilated containers increased *P. japonica* captures by about 100% compared to non-ventilated containers of the same volume. Alm et al. (1996) further showed that capture rates increase significantly when trapped beetles are removed daily, irrespective of container volume. Finally, the small holes drilled at the bottom of the containers to drain excess water likely contributed to reducing insect decomposition rates. Increased container volumes also increase capture rates by preventing them from filling up too quickly under conditions of very high *P. japonica* population densities (>2000-6000 beetles per trap per week) (Alm et al. 1994).

Adding food plant material (grape vine leaves) further increased *P. japonica* capture rates by ∼60% on average, when compared to empty modified traps, or modified traps with only black earth as substrate. Loughrin et al. (1995) also showed that crabapple leaves defoliated by the *P. japonica* produce a complex blend of volatile compounds that serve as reliable cues of a high-quality food source. These results suggest that adding plant material inside traps would then also be useful when using mass trapping techniques to control *P. japonica* populations (e.g., Piñero and Dudenhoeffer 2018).

### Food provision and substrate for the parasitized beetles

Adding substrate or grape vine leaves to the modified trap containers greatly improved *I. aldrichi* development to the puparium stage (61% and 68% pupariation, respectively) compared to empty modified containers (5% pupariation). The addition of a food plant or substrate layer likely helped mimic natural conditions where parasitized beetles bury themselves in the soil just before death, providing the parasitoid with an “air filled chamber” within which it can overwinter (Clausen et al., 1927). These layers probably also help in maintaining a favorable temperature and humidity for the development of immature parasitoids inside traps and helped to prevent the desiccation of trapped insects.

These additions also resulted in increased *I. aldrichi* adult emergence the following season. The addition of both layers inside traps further increased *I. aldrichi* pupariation rates (83%) compared to modified containers with only one layer (61 and 68%). The addition of both layers also resulted in increased parasitoid emergence the following season compared to traps with only a food layer. These results suggest that allowing parasitized *P. japonica* to feed inside the traps is beneficial for *I. aldrichi* larval development and overwintering survival. However, these results were somewhat contradictory with the results of Experiment 2, which indicated that feeding *per se* had a neutral or slightly negative effect on these parameters.

Once collected by hand in the field, we found from Experiment 2 that providing parasitized beetles with food in the lab had a limited impact on the proportion of hosts that yielded puparia. As for other tachinid species, *I. aldrichi* larvae kill its hosts quickly (5-8 days post-oviposition; Clausen et al. 1927; Clausen 1956). Consequently, *I. aldrichi* larvae can be considered as zoonecrophagous (at least during its later developmental stages when the host has died), which may make immature parasitoids less affected by their host’s nutrition than koinobiont species (Dindo, 2011; Dindo & Grenier, 2014). For example, host starvation does not influence the development time of *Compsilura concinnata* larvae (Diptera: Tachinidae) parasitizing the spongy moth (Lepidoptera: Lymantriidae) (Weseloh, 1984), nor pupariation of *Trichopoda giacomellii* (Diptera: Tachinidae) parasitizing *Nezara viridula* (Hemiptera: Pentatomidae) (Coombs, 2004). In Experiment 2, we found that non-fed parasitized beetles yielded slightly more *I. aldrichi* puparia than when food was provided, although this was also associated with reduced weight of the parasitoid puparia. A reduction in *C. concinnata* puparium weight associated with hosts starvation was also reported by Coombs (2004). Based on these results, we suggest that starved beetles are less suitable for *I. aldrichi* larval development. Smaller *I. aldrichi* puparia (i.e., those developing in non-fed hosts) had similar emergence rates to those developing in larger puparia (hosts provided with food). However, tachinid parasitoids emerging from smaller puparia are expected to have reduced size and fecundity (Nakamura, 1995; Allen & Hunt, 2001). Thus, there is a need to balance the disadvantages of providing food to parasitized beetles (higher rearing costs, higher risk of fungal contamination, slightly lower puparium weight) against the advantages of feeding (larger size, longevity and fecundity of emerging adult parasitoids, better capacity to disperse). To better inform this decision, more research is required to explore the magnitude of the observed reduction in puparium size in starved beetles may affect *I. aldrichi* fitness and biological control efficacy.

Previous reports of rearing *I. aldrichi* have not conclusively determined the importance of the rearing substrate for parasitized *P. japonica* (Kidd & McDonald, 1992; Simões & Grenier, 1999). Experiment 2 reported inconsistent but mostly small-magnitude effects of rearing substrate (vermiculite, soil), which may have resulted from differences in the substrates’ water retention and unknown interactions with other factors.

### Recommendations to improve insect collection and parasitoid pupariation

Based on previous knowledge and the results of the present study, we make the following recommendations to optimize collection methods of *P. japonica* parasitized by *I. aldrichi* and rearing the parasitoid to the puparium stage:

1. When small numbers of *I. aldrichi* puparia are needed, we recommend hand-picking parasitized beetles.
2. When large numbers of *I. aldrichi* puparia are needed, we recommend using modified trap containers.
3. Regardless of the collection method, we recommend providing parasitized beetles with plant material. Even though Experiment 2 indicated a slight decrease in pupariation rates with host feeding, we believe it is more advantageous to maximize *I. aldrichi* puparium weight. Increased puparium weight should be translated to adult flies with larger size and fecundity and longevity.
4. Both potting soil and vermiculite are suitable as substrates for rearing *I. aldrichi* to the puparium stage. We recommend vermiculite if puparia need to be retrieved (better color contrast during sorting).
5. Regardless of the collection method, parasitized beetles should be captured early in the season to maximize proportion pupariation (early to late July in the study area).
6. We recommend emptying the traps frequently to decelerate the decomposition of beetles (Alm et al., 1996) and prevent the capture of carrion beetles by-catch (Aubé et al., 2026).
7. When emptying the traps, we recommend rapidly removing carrion beetles by-catch (Aubé et al. 2026) to avoid predation of *I. aldrichi* puparia within the sample.
8. We recommend rearing *I. aldrichi* under protected conditions to avoid potential cases of predation or hyperparasitism (Clausen et al. 1927).

Recommendations on how to store and overwinter *I. aldrichi* puparia and how to manage the emergence of adults are described in Abram et al. (2026).

## Acknowledgements

We thank Nicolas Pomerleau and David Pollender for access to sampling sites and the Jardin Botanique de Montréal for access to overwintering sites. We also thank Marc-Antoine Lampron and Raphaël Boisvert for help in the field and laboratory, and two anonymous reviewers for their useful comments regarding an earlier version of this manuscript.

## Funding

This work was supported by the Natural Sciences and Engineering Research of Canada (NSERC) [grant number 110_2024_2025_Q1_5785] and the Programme Innovation Bioalimentaire 2023-2028, from the Ministère de l’Agriculture, des Pêcheries et de l’Alimentation du Québec (grant number T-08920-B5B0).

## Conflicts of Interest

The authors declare no conflicts of interest.

## Data Availability Statement

The data that support the findings of this study are available from the corresponding author upon reasonable request.

---

**Table S1.** Spring temperature conditions in the incubator in which *Istocheta aldrichi* puparia were held in 2024 (with the previous week’s soil temperature in Agassiz, BC, Canada shown for reference) in Experiment 1.

| Date range | Previous week's soil temperature (°C) | Incubator temperature (°C) |
| --- | --- | --- |
| 21–27 April | 10.4 | 8.4 |
| 28 April–4 May | 10.8 | 9.3 |
| 5–11 May | 12.4 | 10.9 |
| 12–18 May | 14.8 | 13.3 |
| 19–25 May | 13.5 | 12.0 |
| 26 May–1 June | 13.5 | 12.0 |
| 2–8 June | 14.0 | 12.5 |
| 9–15 June | 16.4 | 14.9 |
| 16–22 June | 16.4 | 14.9 |
| 23–29 June | 17.6 | 16.1 |
| 30 June–6 July | 19.2 | 17.7 |
| 7–13 July | 21.5 | 20.0 |
| 14–20 July | 20.9 | 19.4 |
| 21–27 July | 18.9 | 17.4 |
| 28 July–3 August | 18.8 | 17.3 |
| 4–10 August | 19.4 | 17.9 |
| 11–17 August | 19.0 | 17.5 |

## Notes

### Competing Interest Statement

The authors have declared no competing interest.

### Summary of Updates

Minor revisions to main text. Figure 3 separated in two new figures.

## References

Abram PK, Legault S, Doyon J, Makovetski V, Miall JH, Parent J-P, Thiessen J & Brodeur J (2026) Rearing *Istocheta aldrichi* (Diptera: Tachinidae) from field-collected Japanese beetle (*Popillia japonica*): methods to improve overwintering, adult emergence and longevity. bioRxiv. 10.64898/2026.02.10.705140

Abram PK, Parent J-P, Miall JH, Mason PG & Brodeur J (2022) Proposed redistribution of Istocheta aldrichi Mesnil (Diptera: Tachinidae) from Ontario and Québec to British Columbia, Canada, for biological control of the Japanese beetle, Popillia japonica Newman (Coleoptera: Scarabaeidae). https://profils-profiles.science.gc.ca/en/publication/proposed-redistribution-istocheta-aldrichi-mesnil-diptera-tachinidae-ontario-and-quebec

Allen GR & Hunt J (2001) Larval competition, adult fitness, and reproductive strategies in the acoustically orienting ormiine *Homotrixa alleni* (Diptera: Tachinidae). Journal of Insect Behavior 14:283–297.

Alm SR, Yeh T, Campo ML, Dawson CG, Jenkins EB & Simeoni AE (1994) Modified trap designs and heights for increased capture of Japanese beetle adults (Coleoptera: Scarabaeidae). Journal of Economic Entomology 87:775–780.

Alm SR, Yeh T, Dawson CG & Klein MG (1996) Evaluation of trapped beetle repellency, trap height, and string pheromone dispensers on Japanese beetle captures (Coleoptera: Scarabaeidae). Environmental Entomology 25:1274–1278.

Althoff ER & Rice KB (2022) Japanese beetle (Coleoptera: Scarabaeidae) invasion of North America: History, ecology, and management. Journal of Integrated Pest Management 13. 2, 1-11.

Aubé S, Legault S, Doyon J & Brodeur J (2026) Victims of the cure: Determining the pollinator and necrophage biodiversity costs of Japanese beetle [*Popillia japonica*] traps. Biological Conservation 315:111676.

Brodeur J, Doyon J, Abram PK & Parent J-P (2024) *Popillia japonica* Newman, Japanese Beetle / Scarabée japonais (Coleoptera: Scarabaeidae). Biological Control Programmes in Canada, 2013-2023. (ed by MA Vankosky & V Martel) CABI, pp 343–350.

CABI (2021) Classical biological control of Japanese beetle. https://www.cabi.org/projects/classical-biological-control-of-japanese-beetle/

Clausen CP (1956) Biological control of insect pests in the continental United States. United States Department of Agriculture. Technical Bulletin No. 1139, Washington, D.C. USA.

Clausen CP, King JL & Teranishi C (1927) The parasites of *Popillia japonica* in Japan and Chosen (Korea) and their introduction into the United States. United States Department of Agriculture. Department Bulletin No. 1429, Washington, D.C. USA.

Coombs M (2004) Overwintering survival, starvation resistance, and post-diapause reproductive performance of Nezara viridula (L.) (Hemiptera: Pentatomidae) and its parasitoid *Trichopoda giacomellii* Blanchard (Diptera: Tachinidae). Biological Control 30:141–148.

Dindo ML (2011) Tachinid parasitoids: Are they to be considered as koinobionts? BioControl 56:249–255.

Dindo ML & Grenier S (2014) Production of dipteran parasitoids. Mass production of peneficial organisms. (ed by JA Morales-Ramos, MG Rojas & DI Shapiro-Ilan) Elsevier, pp 101–143.

Van Driesche RG (1993) Methods for the field colonization of new biological control agents. T.S Bellows Jr., R.G Van Driesche (Eds.), Proceedings: Thomas Say Publications in Entomology: Steps in Classical Arthropod Biological Control, Entomol. Soc. Am, Lanham (1993), pp. 67–86

Fleming WE (1968) Biological control of the Japanese beetle. Technical bulletin no. 1383. United States Department of Agriculture, Washington, D.C. USA.

Gagnon M-E, Doyon J, Legault S & Brodeur J (2023) The establishment of the association between the Japanese beetle (Coleoptera: Scarabaeidae) and the parasitoid *Istocheta aldrichi* (Diptera: Tachinidae) in Québec, Canada. The Canadian Entomologist 155, e32.

Hopper KR, Roush RT & Powell W (1993) Management of genetics of biological-control introductions. Annual Review of Entomology 38:27–51.

Kidd KA & McDonald RC (1992) Collection and storage of *Istocheta aldrichi* (Mesnil) (Diptera: Tachinidae), a parasitoid of the Japanese beetle, *Popillia japonica* Newman (Coleoptera: Scarabeidae). 1992 Report of Activities. Biological Control Laboratory. Plant Protection Section. North Carolina Department of Agriculture.

King JL (1931) The present status of the established parasites of *Popillia japonica* Newman. Journal of Economic Entomology 24:453–462.

Kreuger B & Potter DA (2001) Diel feeding activity and thermoregulation by Japanese beetles (Coleoptera: Scarabaeidae) within host plant canopies. Environmental Entomology 30:172–180.

Ladd TL & Klein MG (1986) Japanese beetle (Coleoptera: Scarabaeidae) response to color traps baited with phenethyl propionate + eugenol + geraniol (3:7:3) and Japonilure. Journal of Economic Entomology 79:84–86.

Legault S, Doyon J & Brodeur J (2024) Reliability of a commercial trap to estimate population parameters of Japanese beetles, *Popillia japonica*, and parasitism by *Istocheta aldrichi*. Journal of Pest Science 97:575–583.

Lengh R (2021) Emmeans: Estimated marginal means, Aka least-squares means. R package version 1.8. 5.

Leung K, Ras E, Ferguson KB, Ariëns S, Babendreier D, Bijma P, Bourtzis K, Brodeur J, Bruins MA, Centurión A, Chattington SR, Chinchilla-Ramírez M, Dicke M, Fatouros NE, González-Cabrera J, Groot TVM, Haye T, Knapp M, Koskinioti P, Le Hesran S, Lyrakis M, Paspati A, Pérez-Hedo M, Plouvier WN, Schlötterer C, Stahl JM, Thiel A, Urbaneja A, van de Zande L, Verhulst EC, Vet LEM, Visser S, Werren JH, Xia S, Zwaan BJ, Magalhães S, Beukeboom LW & Pannebakker BA (2020) Next-generation biological control: the need for integrating genetics and genomics. Biological Reviews 95:1838–1854.

Loughrin JH, Potter DA & Hamilton-Kemp TR (1995) Volatile compounds induced by herbivory act as aggregation kairomones for the Japanese beetle (*Popillia japonica* Newman). Journal of Chemical Ecology 21:1457–1467.

Ludwig D & Fox H (1938) Growth and survival of Japanese beetle larvae reared in different media. Annals of the Entomological Society of America 31:445–456.

Makovetski V & Abram PK (2024) *Istocheta aldrichi* (Mesnil) makes its biological control debut in British Columbia, Canada. The Tachinid Times 37:4–10.

Makovetski V, Smith ABT & Abram PK (2025) Crowdsourced online data as evidence of absence of non-target attack from the century-old introduction of *Istocheta aldrichi* for biological control of *Popillia japonica* in North America. Journal of Pest Science 98:1451–1462.

McDonald R & Klein M (2023) Establishing the winsome fly, Istocheta aldrichi Mesnil, a major natural enemy of Japanese beetle, Popillia japonica Newman adults.

Nakamura S (1995) Optimal clutch size for maximizing reproductive success in a parasitoid fly, *Exorista japonica* (Diptera: Tachinidae). Applied Entomology and Zoology 30:425–431.

O’Hara JE (2014) New tachinid records for the United States and Canada. The Tachinid Times 27:34–40.

Pelletier M, Legault S, Doyon J & Brodeur J (2023) Where and why do females of the parasitic fly *Istocheta aldrichi* lay their eggs on the body of adult Japanese beetles? Journal of Insect Behavior 36:308–317.

Piñero JC & Dudenhoeffer AP (2018) Mass trapping designs for organic control of the Japanese beetle, *Popillia japonica* (Coleoptera: Scarabaeidae). Pest Management Science 74:1687–1693.

Potter DA & Held DW (2001) Biology and management of the Japanese beetle. Annual Review of Entomology 47:175–216.

Della Rocca F & Milanesi P (2022) The new dominator of the world: Modeling the global distribution of the Japanese beetle under land use and climate change scenarios. Land 11:567.

Sheppard AW, Paynter Q, Mason P, Murphy S, Stoett P, Cowan P, Brodeur J, Warner K, Villegas C, Shaw R & Hinz H (2019) The application of classical biological control for the management of established invasive alien species causing environmental impacts. The Secretariat of the Convention on Biological Diversity. Technical Series No 91. Montreal, Québec, Canada.

Simões AM & Grenier S (1999) An investigation into the overwintering capability of *Istocheta aldrichi* (Mesnil) (Diptera: Tachinidae) a parasitoid of *Popillia japonica* Newman (Coleoptera: Scarabaeidae) on Terceira Island, Azores. Life and Marine Sciences 17A:23–26.

Tumlinson J, Klein M, Doolittle R, Ladd T & Proveaux A (1977) Identification of the female Japanese beetle sex pheromone: Inhibition of male response by an enantiomer. Science 197:789–792.

Vinson SB & Iwantsch GF (1980) Host suitability for insect parasitoids. Annual Review of Entomology 25:397–419.

Vittum PJ (1986) Biology of the Japanese beetle (Coleoptera: Scarabaeidae) in eastern Massachusetts. Journal of Economic Entomology 79:387–391.

Weseloh RM (1984) Effect of size, stress, and ligation of Gypsy moth (Lepidoptera: Lymantriidae) larvae on development of the Tachinid parasite *Compsilura concinnata* Meigen (Diptera: Tachinidae). Annals of the Entomological Society of America 77:423–428.

